# Long-range GABAergic inhibition modulates spatiotemporal dynamics of the output neurons in the olfactory bulb

**DOI:** 10.1101/2020.07.08.194324

**Authors:** Pablo S. Villar, Ruilong Hu, Ricardo C. Araneda

## Abstract

Local interneurons of the olfactory bulb (OB) are densely innervated by long-range GABAergic neurons from the basal forebrain (BF), suggesting that this top-down inhibition regulates early processing in the olfactory system. However, how GABAergic inputs modulate the OB output neurons, the mitral/tufted cells, is unknown. Here, in acute brain slices, we show that optogenetic activation of BF GABAergic inputs produced distinct local circuit effects that can influence the activity of mitral/tufted cells in the spatiotemporal domains. Activation of the GABAergic axons produced a fast disinhibition of mitral/tufted cells consistent with a rapid and synchronous release of GABA onto local interneurons in the glomerular and inframitral circuits of the OB, which also reduced the spike precision of mitral/tufted cells in response to simulated stimuli. In addition, BF GABAergic inhibition modulated local oscillations in a layer-specific manner. The intensity of locally evoked *θ* oscillations was decreased upon activation of top-down inhibition in the glomerular circuit, while evoked *γ* oscillations were reduced by inhibition of granule cells. Furthermore, BF GABAergic input reduced dendrodendritic inhibition in mitral/tufted cells. Together, these results suggest that long-range GABAergic neurons from the BF are well suited to influence temporal and spatial aspects of processing by OB circuits.

## INTRODUCTION

The basal forebrain (BF), a brain region that supports wakefulness, attention and cognition (Anaclet et al., 2015; Xu et al., 2015; Ballinger et al., 2016), has an important role in the state-dependent regulation of sensory circuits (Yang et al., 2014; Hangya et al., 2015; Zant et al., 2016). Among the diverse group of BF neurons, the largest population corresponds to GABAergic projection neurons (Sarter and Bruno, 2002). Yet, unlike the extensive insight on the function of the neighboring BF cholinergic neurons in sensory processing, (Hasselmo, 1995; Linster and Cleland, 2002; Wilson et al., 2004; Parikh and Sarter, 2008; Hellier et al., 2012; Zaborszky et al., 2012; Chapuis and Wilson, 2013; Rothermel et al., 2014) the function of BF GABAergic projections in modulating sensory circuits is not understood. Recent evidence suggests GABAergic neurons provide an important parallel neuromodulatory output from the BF (Gritti et al., 2003; Henny and Jones, 2008; McKenna et al., 2013; Kim et al., 2015; Yang et al., 2017). BF long-range GABAergic neurons (BF-LRGNs) influence the hippocampus and cortex by acting on local inhibitory circuits and modulating the generation of neuronal oscillations, which support essential aspects of the timing of neural activation in these structures (Freund and Antal, 1988; Freund and Meskenaite, 1992; Hangya et al., 2009; Melzer et al., 2012; Gonzalez-Sulser et al., 2014; Kim et al., 2015). Network oscillations are prominent in the OB, the initial site for odor processing, and they are thought to provide a temporal structure for the encoding of odor information (Adrian, 1942; Macrides and Chorover, 1972; Beshel et al., 2007; Schaefer and Margrie, 2007; Junek et al., 2010). The role of local GABAergic neurons in the generation of network rhythms during odor discrimination tasks is well-established (Stopfer et al., 1997; Fukunaga et al., 2014; Osinski and Kay, 2016). In addition, the OB local GABAergic circuits have been involved in decorrelation of principal neurons allowing for discrimination of similar odor (Abraham et al., 2010; Gschwend et al., 2015; Li et al., 2018). Thus, we hypothesize that by modulating local inhibitory circuits BF-LRGNs could influence odor processing in the OB. In agreement with this possibility, chemogenetic silencing of LRGNs of the magnocellular preoptic nucleus (MCPO), a main source of BF GABAergic inhibition to the OB (Gracia-Llanes et al., 2010), produces a notable reduction in the discrimination of similar odors (Nunez-Parra et al., 2013); however, how this BF inhibition influences neuronal circuits in the OB remains unclear.

Here, we used a combination of conditional genetics, immunohistochemistry and electrophysiology in acute brain slices to define the physiological framework by which the BF GABAergic projections modulate, at the circuit level, the spatiotemporal dynamics of the output neurons in the OB. We first established that MCPO Gad2 neurons, which comprise the main inhibitory projections to the OB, appear phenotypically homogeneous, using GABA as the main transmitter, unlike other GABAergic neurons in the BF (Saunders et al., 2015; Case et al., 2017). In agreement with previous work (Gracia-Llanes et al., 2010), immunohistochemical analysis revealed that BF-LRGNs extensively innervate the granule cell layer (GCL) and to a lesser extent the glomerular layer. Consistent with this anatomical distribution, phasic activation of BF GABAergic axons, elicited fast inhibitory responses in local inhibitory neurons, including the granule cells (GCs) and periglomerular cells (PGCs); however, inhibitory responses were absent in the output neurons, the mitral and tufted cells (MC and TCs, respectively). Functionally, the selective activation of the GABAergic axons in the OB results in a disinhibitory effect of the output neurons; activation BF inhibition increased the firing rate of active MCs. We show that this increase in firing rate can result from a reduction in the inhibition by glomerular inhibitory neurons and by a reduction in dendrodendritic inhibition at GC-MC synapses. In addition, top-down inhibition decreased the spike precision of MCs in response to simulated sensory stimuli. Importantly, activation of BF GABAergic inputs produced a significant reduction in the power of local *θ* and *γ* oscillations, thus desynchronizing the rhythmic activity in the OB. Together, these results indicate that fast BF GABAergic inhibition is well suited to modulate early stages of odor processing by regulating spatiotemporal dynamics of MCs.

## RESULTS

### GABAergic neurons in the MCPO innervate inhibitory circuits of the OB

Previous studies have shown that OB projecting LRGNs are clustered in a lateral region of the BF, the MCPO (Gracia-Llanes et al., 2010). To broadly label these projections neurons we used Gad2-Cre mice, as the GABAergic marker Gad2 is abundantly expressed in the MCPO (Nunez-Parra et al., 2013). Gad2-Cre mice were injected with the anterograde virus AAV5-Flex-tdTomato in the MCPO (**Figure 1A**, diagram). In agreement with previous work (Gracia-Llanes et al., 2010; Nunez-Parra et al., 2013), this anterograde injection resulted in extensive labelling of fibers in the OB, with a distinct pattern of labelling across its cellular layers. Fluorescently labeled axons of LRGNs exhibited a non-uniform distribution pattern throughout all the layers of the OB, with dense labelling in the GCL and to a lower extent in the external plexiform layer (EPL) (**Figure 1A**). Similarly, there was significant innervation around juxtaglomerular neurons in the glomerular layer (GL) (mean normalized pixel intensity GL, 0.35 ± 0.07; EPL, 0.15 ± 0.02; GCL, 0.7 ± 0.09; n= 6 slices, 3 mice). At a cellular level, the GABAergic axons were characterized by thick and smooth processes running along the distinct cellular layers, with profuse ramifications and axonal boutons (**Figure 1B**, arrow heads).

**Figure 1.**
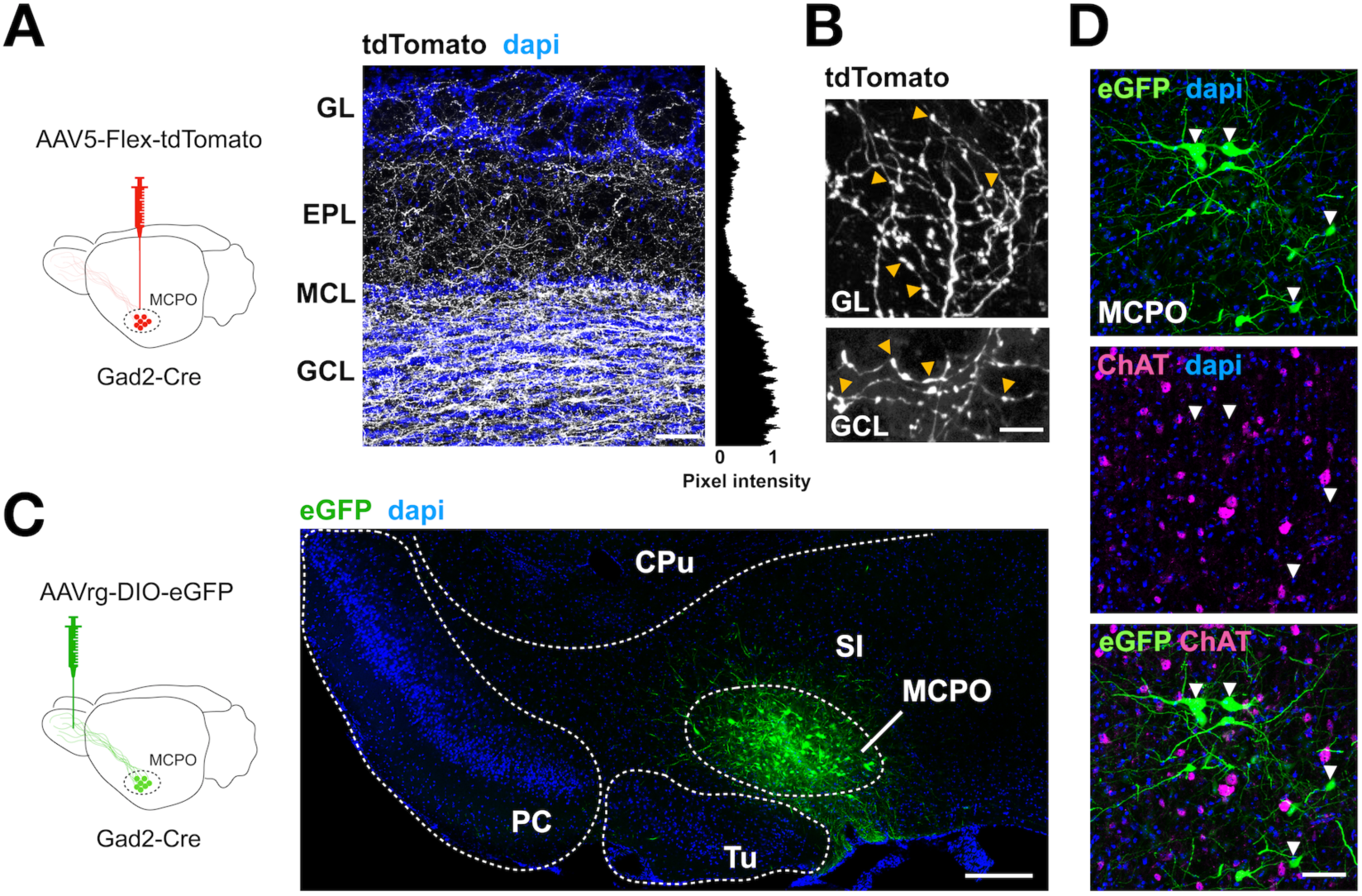
GABAergic projection neurons to the OB are clustered in the MCPO region of the BF and are different from cholinergic neurons. **(A)** Left, diagram of the anterograde approach to label MCPO GABAergic neurons; a AAV5-Flex-tdTomato virus was injected in the MCPO of Gad2-Cre mice. Right, confocal image of a section of the main olfactory bulb (MOB; n= 6) showing the distribution pattern of Gad2-tdTomato axons (shown in white, to enhance the contrast of the staining) across the different cell layers, revealed by the nuclear dye dapi (blue). The mean normalized pixel intensity across layers is shown on the right. The densest distribution of Gad2 axons is found in the granule cell layer (GCL) and the glomerular layer (GL) of the MOB. MCL, mitral cell layer; EPL, external plexiform layer. The scale bar is 100 μm. **(B)** MCPO GABAergic axons innervating the GL (top) and GCL (bottom) exhibit numerous boutons (yellow arrowheads). The scale bar is 10 μm. **(C)** Left, diagram of the approach to retrogradely label the MCPO GABAergic neurons; a AAVrg-DIO-eGFP virus was injected unilaterally in the OB of Gad2-Cre mice. Right, confocal micrograph showing that transduced Gad2-eGFP positive neurons (green) are clustered in the MCPO (CPu, caudate putamen; PC, piriform cortex; Tu, olfactory tubercle; SI, substantia innominata; MCPO, magnocellular preoptic area). Scale bar: 1 mm. **(D)** High magnification confocal micrographs of the MCPO containing GFP transduced Gad2 neurons (green), immunostained with antibody against the cholinergic marker ChAT (magenta). Several neurons are positive for ChAT in the MCPO region; however, this representative image illustrates the lack of colocalization of the cholinergic marker and the GABAergic neurons retrolabeled from the OB (white arrowheads). The scale bar is 40 μm.

To specifically access the population of OB-projecting GABAergic neurons in the MCPO we used an AAV variant that produces efficient retrograde labeling, the rAAV2-retro virus (Tervo et al., 2016; in ‘t Zandt et al., 2019). This virus was injected unilaterally into the OB of Gad2-Cre mice (**Figure 1C**, diagram). Approximately three weeks after injection, transduced GABAergic neurons were abundant in the ipsilateral hemisphere and confined to the MCPO (**Figure 1C**). Recent work has shown that in the medial septum/diagonal band of Broca axis (MS/DBB), a subregion of the BF, some neurons express both GABAergic and cholinergic markers, suggesting that MCPO GABAergic neurons could exhibit a mixed phenotype (Saunders et al., 2015; Takács et al., 2018). To evaluate this possibility, we immuno-stained retrolabeled GABAergic MCPO neurons with an antibody directed against the enzyme choline acetyl transferase (ChAT), a cholinergic marker. As shown in **Figure 1D**, several putative cholinergic neurons are labelled in this region, however, we observed minimal labelling of ChAT protein among retrolabeled MCPO GABAergic neurons, with sections showing only a ∼1 % of colocalization (GFP+= 832 neurons; ChAT+= 468; GFP+/ChAT+= 10; n= 12 slices, 3 mice). This observation is in agreement with a recent study showing absence of colocalization of OB projecting BF neurons with ChAT (Hanson et al., 2020). In contrast, when Cre-dependent expression of the fluorescent protein eGFP was achieved by direct transduction in the MCPO, 16.5% of Gad2 positive (Gad2+) neurons displayed colocalization with the cholinergic marker, as previously reported (Saunders et al., 2015; Sanz Diez et al., 2019) (GFP+= 510 neurons; ChAT+= 423; GFP+/ChAT+= 84; n= 9 slices, 3 mice). Thus, although we cannot rule out the possibility of low levels of expression of ChAT that were undetected by our immunoassay, or the presence of other neurotransmitters released by Gad2+ neurons, our data indicates that MCPO GABAergic neurons that project to the OB exhibit mostly a GABAergic phenotype (see also **Figure 3C**).

**Figure 2.**
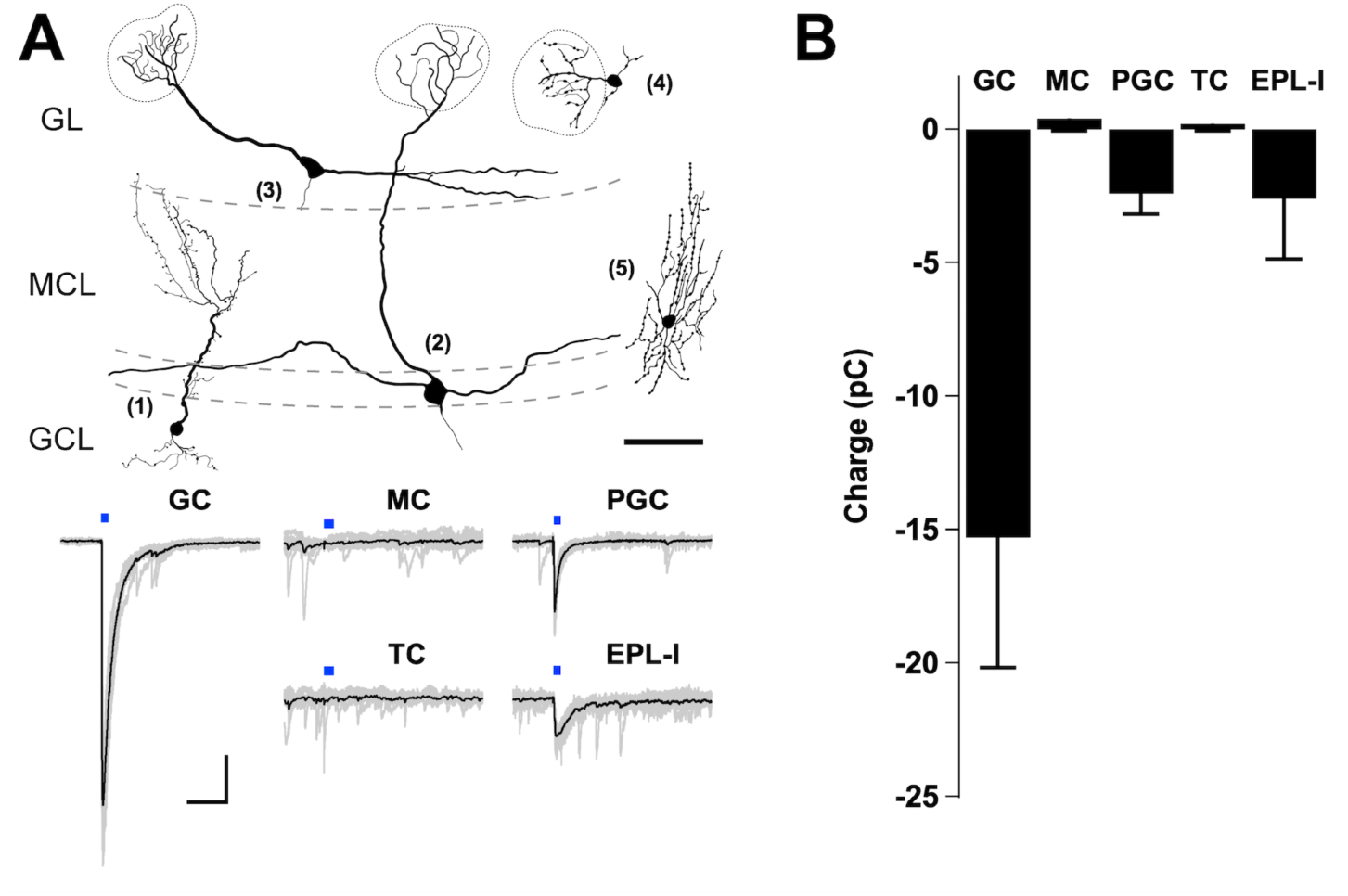
Inhibitory neurons are postsynaptic partners of MCPO long-range GABAergic neurons in the OB. **(A)** Upper drawings, example of distinct reconstructed neurons, post-recording; 1, granule cell (GC); 2, mitral cell (MC); 3, tufted cell (TC); 4, periglomerular cell (PGC): 5, external plexiform layer interneuron (EPL-I). The morphology of the neurons was reconstructed from confocal images of fixed cells that were filled with Alexa Fluor-594 during the recordings. The scale bar is 100 μm. Bottom, example of eIPSCs recorded at −70 mV in symmetrical chloride conditions, upon stimulation of GABAergic axons expressing ChR2 with blue light (5 ms). LED stimulation elicited large inward currents in GC, PGC and EPL-I but not in output neurons, the MC or TC. The scale bar is 200 ms and 50 pA **(B)** Bar graph showing the total charge transferred during the GABAergic eIPSCs in distinct cell types in the OB (GC, n= 16; MC, n= 11; PGC, n= 5; TC, n= 10; EPL-I, n= 3). Responses are observed in the main inhibitory types, but not in the output neurons.

**Figure 3.**
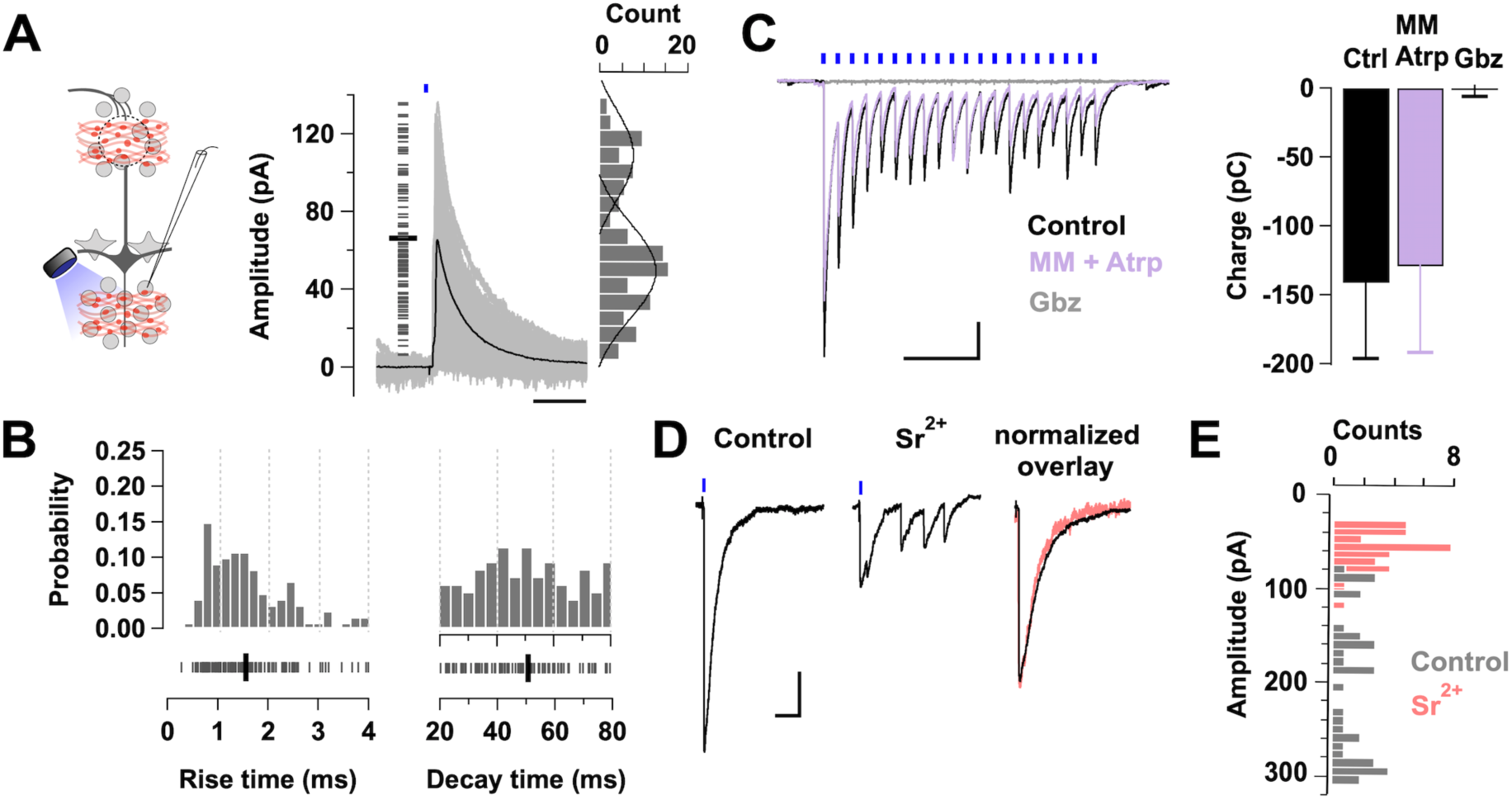
Synaptic properties of the BF long-range GABAergic inputs onto GCs. **(A)** Left, diagram of the experimental configuration; GCs were recorded in voltage clamp, while GABAergic axons expressing ChR2 were stimulated with a brief pulse of light (0.5-1 ms). Right, overlay of selected inhibitory postsynaptic currents evoked with minimal stimulation (min-eIPSCs, gray traces) in GCs (n= 5 cells, 3 mice). Only min-eIPSCs with rise times of less than 4 ms are included (n= 120 events, 5 cells). The short gray lines on the left correspond to the amplitude of single events. The min-eIPSC had an average amplitude of 66 ± 3 pA (thick black line). The amplitude histogram of the min-eIPSCs is shown on the right. Amplitudes show a bimodal distribution with a small peak centered at 48 pA and a higher peak at 100 pA. The black lines correspond to the fitting of two gaussian distributions to the amplitude distribution. The scale bar is 100 ms. **(B)** Probability distribution histograms for the rise time (left, 10-90% of the peak) and decay time (right, τ_w_) of the min-eIPSC events shown in A. An equivalent number of events were taken from each cell (median 26). The ticks at the bottom correspond to the values for each event. The average rise time (1.6 ± 0.1 ms) and decay time (50.6 ± 1.8 ms) are shown by thick black lines. **(C)** Evoked IPSCs (eIPSCs) recorded in GCs, using a CsCl based internal solution, in response to LED stimulation (5 ms, 10 Hz). At this frequency of stimulation, the peak decreases in time but each eIPSC appear synchronous throughout the train (black traces). The eIPSCs are unaffected by the perfusion of a mixture of the cholinergic blockers mecamylamine (MM, 10 µM) and atropine (Atrp, 3 µM) (purple trace; n= 5, p= 0.25), but completely blocked by the GABA_A_R blocker gabazine (Gbz, 10 µM) (gray trace; n= 3, p= 0.04). The scale bar is 500 ms and 30 pA. **(D)** Left, light evoked IPSCs in GCs are desynchronized by the equimolar replacement of calcium by strontium (Sr^+2^, 2 mM). Right, overlay of peak-normalized IPSCs for control (black) and strontium (pink) showing similar kinetics. The scale bar is 200 ms and 50 pA. The holding potential in C and D is −70 mV. **(E)** Histogram overlaying the eIPSC amplitudes during control (gray) and strontium (pink) application (n= 3 cells)

### Activation of LRGN produces a fast inhibition in local inhibitory neurons of the OB

We next examined the influence of endogenously released GABA from BF GABAergic axons in the most prominent components of the OB circuit. To selectively activate GABA release from GABAergic axons in the MOB, we expressed the light-gated cation channel channelrhodopsin-2 (ChR2) in the MCPO of Gad2-Cre mice and conducted targeted recordings from different cell types across the OB (**Figure 2**). We maximized the probability of detecting evoked GABA currents, by performing these recordings in symmetrical chloride conditions (see Methods), in which GABA elicits large inward currents. Light stimulation, reliably evoked short latency inhibitory postsynaptic currents (eIPSCs) in two of the most prominent inhibitory neurons of the MOB; the GCs and the PGCs (**Figure 2B;** onset: GCs, 6.8 ± 0.7 ms, n= 14; PGCs, 7.2 ± 1 ms, n= 5). The amplitude and kinetics of the currents was variable among these different cell types. Quantification of the transferred charge (see Methods) indicated that average inhibitory responses were significantly larger in the GCs (**Figure 2B**, GCs, −15.3 ± 5 pC, n= 16 vs PGCs, −2.4 ± 0.8 pC, n= 5; p= 0.02), and that GCs also exhibited eIPSCs with a slower decay time (GCs, 60.3 ± 7.2, ms n= 16 vs. PGCs, 31.7 ± 4.8 ms, n= 5, p= 0.04). Additionally, short-latency GABAergic responses were also observed in EPL medium-sized interneurons (EPL-I), which presumably correspond to the fast-spiking (FS) interneurons described in this region (EPL-I, −2.6 ± 2.2 pC, n= 3, **Figure 2A, B**) (Hamilton et al., 2005; Huang et al., 2013). In contrast, light stimulation failed to produce any detectable inhibitory current in the output neurons of the OB, the MCs and TCs (MCs, −0.4 ± 0.5 pC, n= 11; TCs, 0.2 ± 0.3 pC, n= 10). These results are consistent with a recent report that examined the targets of GABAergic neurons from a different region of the basal forebrain (Hanson et al., 2020). Additionally, BF GABAergic axons produced a similar pattern of labelling in the accessory OB (AOB), a region involved in pheromonal signal processing, with dense innervation of the GCL (**Figure S1, A**). Similar to the main OB, GABA release from MCPO axons elicited eIPSCs only in the inhibitory cell types of the AOB (**Figure S1 B-C**; GCs, −11 ± 4 pC, n= 12; PGCs, −5 ± 3.8 pC, n= 5; M/TCs, −0.9 ± 0.7 pC, n= 6). Together, these results indicate that, at the circuit level, BF inhibition functionally targets inhibitory but not excitatory neurons in the OB.

### Synaptic activation of GCs by BF-LRGN input is synchronous and long-lasting

To further determine the impact of the BF inhibition onto the local inhibitory neurons, we examined the synaptic properties of MCPO inhibitory inputs onto GCs, which showed the densest innervation by LRGNs. GABA release in the OB was evoked from LRGN axon terminals expressing ChR2 by a brief light stimulation pulse (0.5-1 ms). The duration of the light stimulation was adjusted to reduce the probability of stimulating multiple axons simultaneously (achieving a ∼40% failure rate). We termed this a minimally evoked IPSC (min-eIPSC) (Banks et al., 1998; Hagiwara et al., 2012). We recorded the min-eIPSC at 0 mV, using a Cs-gluconate based internal solution (see Methods), which allowed us to isolate the outward GABAergic currents, without affecting the function of local circuits by the use of synaptic transmission blockers. Light stimulation elicited a short latency min-eIPSC (mean ± SD, 8.1 ± 2.8 ms, n= 5 cells), which occurred with a variable onset likely due to differences in axonal geometry and the short duration of the stimulation (**Figure 3A**). The average amplitude of the min-eIPSC was 66 ± 3 pA (**Figure 3A**), with kinetics characterized by a fast rise time (10-90 % of the peak, 1.6 ± 0.1 ms) and a slower decay time (50.6 ± 1.8 ms) (**Figure 3B**). The min-eIPSC amplitudes exhibited a bimodal distribution, having a small amplitude peak (mean ± SD, 48 ± 15 pA) and a larger peak (mean ± SD, 100 ± 31 pA). The majority of the events had fast rise times (75% <2 ms), which included the majority of the larger amplitude events, with a smaller number of events (∼25%) having slower rise times. The events of larger amplitude and faster rise time likely reflect a predominant perisomatic targeting of the MCPO input onto GCs, while the smaller amplitude and slower rise time event reflecting more distal GABAergic inputs (**Figure 1A, Figure 3B**, left). In contrast, the decay times showed a monophasic distribution due to their longer time course and thus subjected to less apparent filtering (**Figure 3B**, right). Interestingly, the decay time of the evoked min-IPSC from MCPO axons is relatively slow compared to the IPSCs driven in GCs by local GABAergic neurons such as the deep short-axon cells (dSAC, τ∼10 ms) (Eyre et al., 2008). The min-eIPSC decay time in GCs is also slower than the decay time of spontaneous IPSCs from GC activity recorded in MCs under similar conditions (23 ± 0.7 ms, n= 51, not shown). The relatively slower current relaxation of the BF GABAergic inputs suggests they have a longer temporal influence in GCs compared with the influence of local inhibition. Importantly, while the light-elicited currents were completely abolished by the GABA_A_R blocker gabazine (Gbz; control −89 ± 21 pC vs. Gbz, 0.3 ± 1.5 pC, n= 3, p= 0.04), they were unaffected by a mixture of the cholinergic receptor blockers, mecamylamine (MM) and atropine (Atrp) (control −141 ± 55 pC vs. cholinergic blockers, −130 ± 62, p= 0.25, n= 5), further indicating that MCPO inputs onto GCs are mostly GABAergic (**Figure 3C**).

Synchronized vesicular release is a common feature of evoked neurotransmission in the nervous system and accounts for phasic synaptic transmission, while asynchronous release provides persistent neurotransmitter release favoring delayed transmission (Atluri and Regehr, 1998; Hefft and Jonas, 2005; Südhof, 2013; Wen et al., 2013; Kaeser and Regehr, 2014). Our data indicates that MCPO inputs produce a fast-synchronized release of GABA onto GCs, as light stimulation of MCPO axons with a single pulse, or across a high frequency stimulation train, always evoked currents that decay monotonically (**Figure 3A, C, D)**. Accordingly, equimolar replacement of the extracellular Ca^2+^ by Sr^2+^, a divalent ion that disrupts synchronized vesicular release (Dodge et al., 1969; Goda and Stevens, 1994; Xu-Friedman and Regehr, 2000; Shin et al., 2003), resulted in a barrage of smaller current events upon light stimulation (**Figure 3D**, left). In normal Ca^2+^, the eIPSC had a mean amplitude of 191 ± 12 pA, while in the presence of Sr^2+^ the current amplitude was significantly lower (mean 54 ± 4 pA, p<0.001, n= 3) (**Figure 3E**). Importantly, in agreement with a quantal mechanism of release at these synapses, the kinetics of eIPSC evoked in the presence of Sr^2+^ closely resembled the kinetics of those in normal Ca^2+^ (**Figure 3D**, right) (rise time, in Ca^2+^, 1.52 ± 0.3 ms vs. in Sr^2+^, 1.85 ± 0.1 ms, p= 0.3; decay time, in Ca^2+^, 63.6 ± 2 ms vs. in Sr^2+^, 60.5 ± 3 ms, p= 0.43, n= 3). Together these results suggest that activation of BF-LRGNs axons in the OB produces a fast-synchronous release of GABA onto GCs, likely from multiple synaptic contacts.

### Activation of BF-LRGNs disinhibit mitral cells and modulates the extent of lateral inhibition

We next examined the influence of the fast and synchronous GABA release onto the local inhibitory networks of the OB by BF-LRGN. In the glomeruli, local GABAergic neurons drive feedforward inhibition onto MCs, providing a mechanism for sensory gain control and decorrelation of odor representations (Wilson and Mainen, 2006; Zhu et al., 2013; Banerjee et al., 2015). Thus, inhibition of PGCs by BF-LRGNs (**Figure 2**) suggests that BF inhibition can modulate the extent of feedforward inhibition in glomerular domains. To examine this possibility, we evoked activity in MCs by stimulating the axons of OSNs in the olfactory nerve (ON), while locally stimulating GABA release from BF-LRGNs axons in the GL (**Figure 4A**, diagram). Stimulation of the ON produced a long-lasting depolarization in MCs, with sustained firing of action potentials (**Figure 4A**), which was significantly increased by simultaneous light stimulation of BF-LRGNs axons, in agreement with the disinhibitory effect on this afferent input (firing rate, control: 4.9 ± 5.1 Hz, + LED: 8 ± 3.4 Hz, n= 2). We next recorded simultaneously excitatory and inhibitory currents in MCs evoked by ON stimulation, using symmetrical chloride, at a holding potential of −60 mV (**Figure 4B**). The ON stimulation evoked a large inward current consisting of a barrage of glutamatergic and GABAergic events which was reduced ∼60 % by light stimulation directed to the GL (control, −137 ± 20 pC; +LED, −53 ± 15 pC; n= 10, p<0.001) (**Figure 4C**). In agreement with the possibility that the reduction in the inward current by light stimulation was mostly due to a reduction of the GABAergic component, blockade of GABA_A_ Rs produced a similar reduction (∼57 %) in the current evoked by ON stimulation (control: −182 ± 30 pC, Gbz: −73.5 ± 11 pC, n= 4, p= 0.003). Lastly, the ON evoked current was completely abolished in the presence of gabazine and the broad glutamate receptor blocker kynurenic acid (1 mM) (control: −142 ± 32 pC, + blockers: −6.7 ± 4.5 pC, n= 6, p= 0.01). These results are consistent with an inhibitory control of LRGNs on glomerular GABAergic neurons targeting MCs.

**Figure 4.**
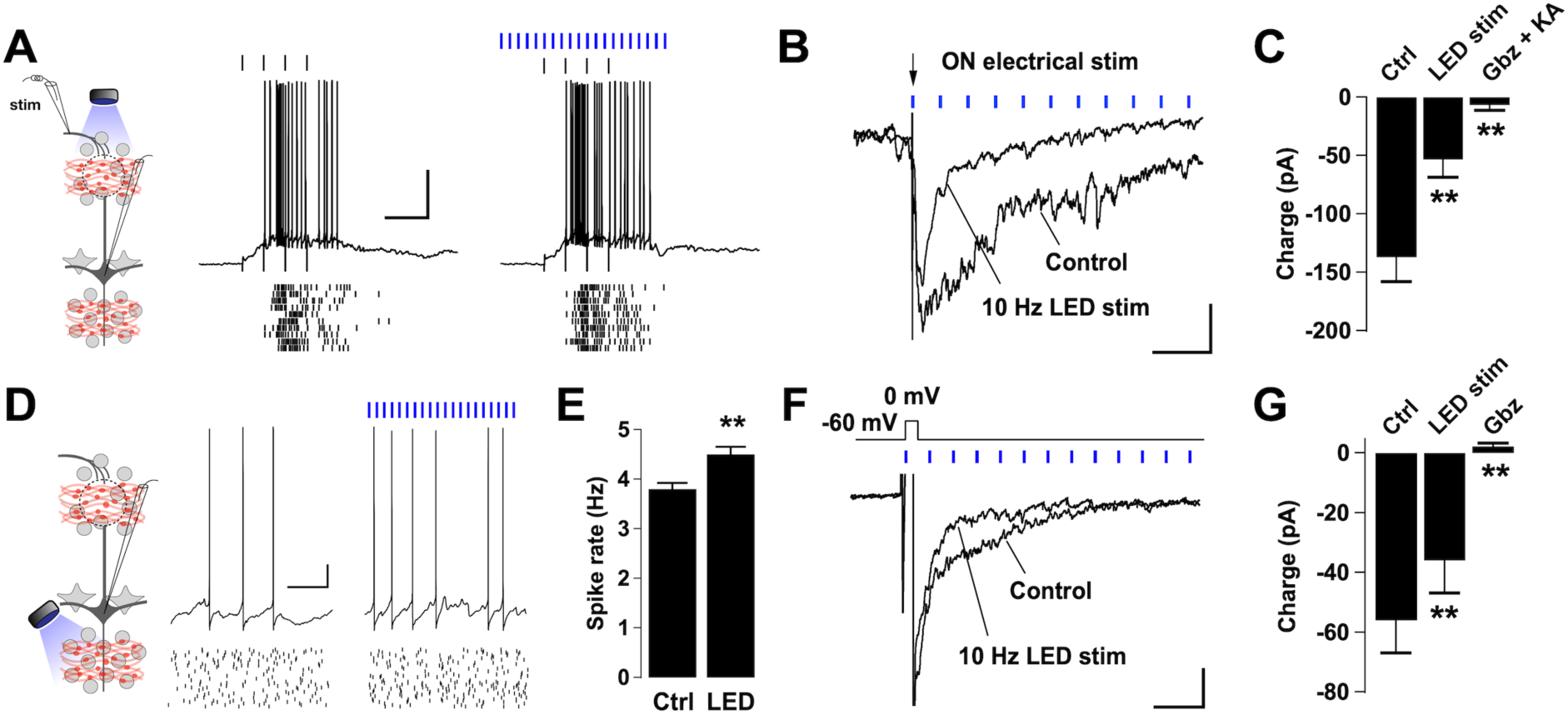
Activation of BF GABAergic inputs disinhibits mitral cells and reduces dendrodendritic inhibition. **(A)** Left, diagram of the experimental configuration; MCs where recorded either in current or voltage clamp while the sensory axons in the olfactory nerve (ON) were activated by electrical stimulation. BF GABAergic axons expressing ChR2 were activated by blue light in the GL. Right, responses in a representative MC recorded while stimulating the ON with a glass electrode (100 μA, 100 μs, 4 Hz, top black ticks) in the presence (left) or absence (right) of LED stimulation (at 10 Hz, blue ticks). The stimulus intensity was adjusted to elicit firing in the MC. Bottom, spike raster plots for 10 trials in the cell shown above. The membrane potential was −57 mV (with zero current injection). **(B)** Synaptic currents evoked in a MC by electrical stimulation of the ON (100 μA, 100 μs, arrow). Recording were performed in symmetrical chloride, in which excitatory and inhibitory currents are seen as inward deflections. A single ON stimulation produced a long-lasting inward current, which was reduced in the presence of LED stimulation in the GL (blue ticks). The holding potential is −60 mV. The scale bar is 200 ms and 50 pA. **(C)** The large barrage of evoked synaptic activity by the ON stimulation is greatly suppressed by LED stimulation (n= 10, p<0.001), and completely abolished by blockers of GABA_A_ and glutamate receptors (Gbz, 10 μM; kynurenic acid, KA, 1 mM, respectively; n= 6, p= 0.01). **(D)** Left, diagram of the experimental arrangement; MCs were recorded either in current or voltage clamp while LED stimulation was directed to the GCL. Right, voltage traces of a representative MC held at peri-threshold membrane potential in control and in the presence of LED stimulation (4 Hz). Spike raster plots for 20 trials are shown in the traces below. **(E)** Summary bar graphs for spike frequency showing a significant increase in the firing rate during the LED stimulation compared to control (n= 6, p= 0.004). **(F)** Overlaid of average current traces showing dendrodendritic inhibition on a MC evoked by a short depolarization (0 mV, 50 ms) in control and in the presence of LED stimulation (10 Hz). The scale bar is 200 ms and 100 pA; the holding potential is −60 mV. **(G)** Summary bar plot showing a significant difference in the synaptic charge transferred in control versus during LED stimulation (n= 8, p= 0.003). Consistently, Gbz (10 μM) completely blocked the evoked dendrodendritic current in MCs (n= 10, p= 0.0002).

Dendrodendritic inhibition (DDI) at MC-GC synapses is thought to shape the output signal of MCs both in the temporal and spatial domains, through recurrent and lateral inhibition (Yokoi et al., 1995; Isaacson and Strowbridge, 1998; Christie et al., 2001; Shepherd, 2004). Therefore, we next examined how BF inhibition shapes the responses of MCs, by locally stimulating GABA release from MCPO axons in the GCL. We depolarized MCs by constant current injection to produce a low firing rate (∼4Hz, **Figure 4D**). In agreement with a disinhibitory action of the BF afferent input, via inhibition of GCs, light stimulation directed to the GCL significantly increased the basal firing rate in MCs (control, 3.8 ± 2.3 Hz; +LED, 4.5 ± 2.5 Hz; n= 6, p= 0.004) (**Figure 4E**). In order to directly examine the modulation of DDI by BF inhibition, we recorded the GABAergic currents evoked by a depolarizing pulse in MCs while holding the cells at −60 mV (**Figure 4F**). A brief stimulation (50 ms) elicited a barrage of GABAergic currents with a relaxation time of 640 ± 230 ms (n= 8) similar to the values previously described (Isaacson and Strowbridge, 1998; Schoppa, 1998). In the presence of local stimulation of GABA release from BF-LRGNs axons, the DDI was significantly reduced (**Figure 4F-G**) (control, −56 ± 11 pC, LED stimulation, −36 ± 11 pC, n= 8, p= 0.003). As expected, blocking GABA_A_Rs completely abolished the evoked DDI in MCs (control, −52 ± 14 pC; Gbz, 2 ± 1 pC; n= 10, p<0.001). These results suggest that activation of BF-LRGNs can reduce the extent of DDI in MCs and thus can influence odor processing by reducing lateral inhibition.

### BF-LRGNs modulate *θ* and *γ* oscillations in a layer specific manner

Inhibition from local GABAergic circuits contributes to generate a temporal framework in which low and high frequency neuronal oscillations exist in the OB (Kay et al., 2009; Wachowiak, 2011). Although the underlying mechanism is not completely understood, oscillations in the *θ* frequency band (2-12 Hz) entrained by the respiratory cycle, are orchestrated by PGCs (Lagier et al., 2004; Fukunaga et al., 2014), while *γ* oscillations (25-85 Hz) require the activation of GCs (Rall and Shepherd, 1968; Balu et al., 2007; Lagier et al., 2007; Kay, 2014). Since a main target of BF inhibition are the PGCs and GCs, we hypothesized that BF GABAergic inhibition could differentially influence the generation of oscillatory activity in the OB by regulating the activity of the glomerular and infra-mitral inhibitory circuits. To examine this possibility, we recorded the local field potentials (LFPs) evoked by stimulation of the ON (Lagier et al., 2007), while optogenetically inducing GABA release from MCPO GABAergic axons (**Figure 5A**, diagram). A brief, high frequency, electrical stimulation of the ON (100 Hz, 50 ms) elicited both slow and fast fluctuations in the LFP, that persisted for ∼1 s following the cessation of the ON stimulation (**Figure 5B-C**). Frequency analysis of the LFP signals revealed that both *θ* and *γ* oscillations concurred; they were apparent in both the raw and filtered LFP traces (**Figure 5B-D)**. Importantly, when the MCPO GABAergic axons were locally stimulated in the GL, the power of *θ* was significantly reduced (*θ*: control 1.56 ± 0.3, LED 1.02 ± 0.2, n= 5, p= 0.05). We also observed a trend towards a lower *γ* power, albeit this was not significant (*γ*: control 1.36 ± 0.2, LED 1.1 ± 0.1, n= 5, p= 0.09) (**Figure 5E**, right). In contrast, when the light stimulation was directed to the GCL, the power of *γ*, but not θ, was significantly reduced (θ: control 1.24 ± 0.2, LED 1.18 ± 0.2, n= 5, p= 0.66; *γ*: control 1.24 ± 0.1, LED 1.11 ± 0.1, n= 5, p= 0.04) (**Figure 5E**, left). These results suggest that the BF-LRGNs could differentially regulate the dynamics of local GABAergic circuits in the GL and GCL. We note that LED stimulation alone, in either GL or GCL, failed to induce significant changes in the LFP, owing perhaps to the low activity of inhibitory circuits in the slice. Nevertheless, a mixture of the excitatory and inhibitory synaptic blockers, kynurenic acid and Gbz, completely abolished the electrically induced oscillations, in agreement with their synaptic origin (data not shown).

**Figure 5.**
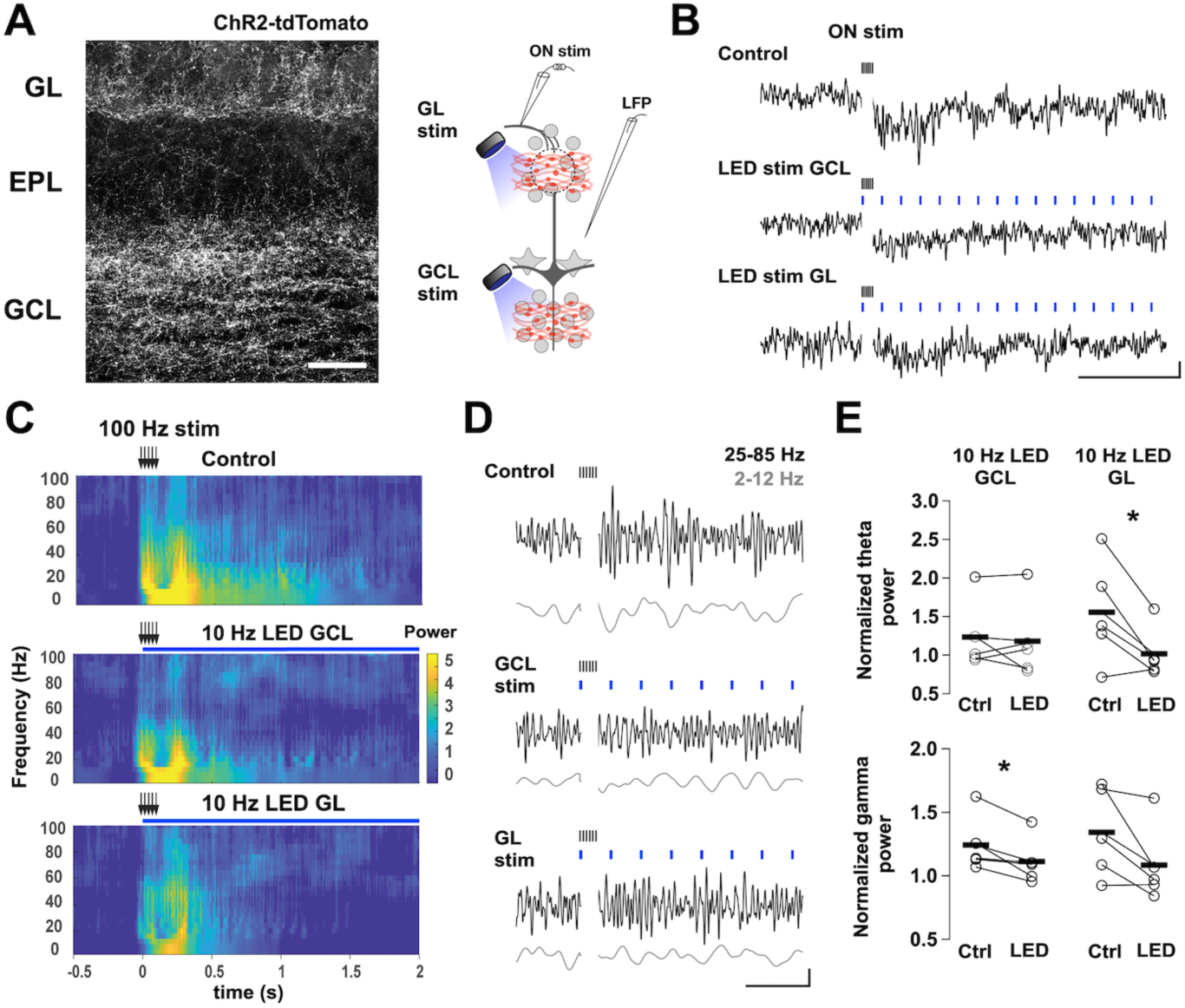
Layer specific modulation of local field potential oscillations by activation of BF GABAergic inputs. **(A)** Left, image of a recorded section of OB showing expression of ChR2-tdTomato achieved by an injection of AAV5-Flex-ChR2-tdTomato virus in the MCPO. Right, diagram of the experimental configuration; a low resistance patch electrode was placed in the external plexiform layer (EPL) to record the local field potential (LFP) in OB slices containing BF GABAergic axons expressing ChR2. Oscillatory activity was elicited by stimulating the olfactory nerve (ON) with a brief high frequency stimulus (100 μA, 100 Hz for 50 ms). In alternate trials, we stimulated the BF GABAergic axons with a blue LED (5 ms, 10 Hz for 2 s) directed to the GCL or the GL using a 40x objective (focused ∼400 μm apart). The scale bar is 100 μm. **(B)** Sample traces of LFP recordings in the EPL during electrical stimulation of the ON (black ticks) in control (upper trace) and with LED stimulation over the GCL (blue ticks, middle trace) or the GL (bottom trace). The scale bar is 20 μV and 500 ms. **(C)** Power spectra of the LFP recorded under the different conditions as in B. Power was normalized respect to the pre stimulation period. The power spectra show significant activity in the 2-50 Hz oscillatory range, which include both *θ* and *γ* bands, elicited by the ON stimulus. Light activation of BF GABAergic axons in the GCL and GL modulate the power in this frequency range. **(D)** Band pass filtered LFP traces for the different conditions; low frequency, 2-12 Hz (*θ*, grey), and high frequency, 25-85 Hz (*γ*, black). The scale bar is 10 μV and 200 ms. **(E)** Pair comparison of the normalized power of the *θ* (upper plots) and *γ* frequency bands (lower plots) in the absence (control) and presence of light stimulation (LED). Light stimulation in the GL significantly reduced the power of the *θ* band (n= 5, p= 0.05), but not the *γ* band (n= 5, p= 0.09), while LED stimulation in the GCL significantly reduced the power of the *γ* band (n= 5, p= 0.04), but not the *θ* band (n=5, p= 0.66).

### Activation of BF-LRGNs inputs decreases spike precision in mitral cells

In other brain regions, long-range GABAergic inhibition influences rhythmic activity through direct modulation of local GABAergic interneurons, which in turn can regulate the precision of firing of principal neurons (Tamamaki and Tomioka, 2010; Melzer et al., 2012; Kim et al., 2015). In the MOB spike precision in MCs can be regulated by inhibition from GCs (Schoppa, 2006), a main target of BF inhibition; therefore, we next examined how BF inhibition modulates spike precision of MCs. We simulated the occurrence of coincident sensory inputs of increasing synchrony onto MCs, overlaid on a 4 Hz respiration-like wave. The currents that produced the simulated excitatory postsynaptic potentials (sim-EPSPs) were adjusted to elicit a similar firing rate across trials (**Figure 6A**, top current trace) and the first 4 and last 4 stimuli were averaged to represent lower and higher synchrony, respectively. Under control conditions, the jitter in the spiking generated by the low synchrony sim-EPSCs was higher, compared to the high synchrony sim-EPSCs; in other words, the spike precision is higher with the high synchrony stimuli (Rodriguez-Molina et al., 2007) (**Figure 6A**, lower traces; low synchrony 34 ± 2.3 ms vs. high synchrony 18.9 ± 2.2 ms; n= 7, p= 0.02). As expected, the overall firing rate of MCs during the sim-EPSPs significantly increased when the MCPO GABAergic axons were locally stimulated with light (**Figure 6B**; control 7.2 ± 1.1 vs. + LED 8.3 ± 1.3 Hz, n= 7, p= 0.02). Importantly, optogenetic activation of BF-LRGNs axons significantly increased spike jitter in MCs for sim-EPSCs at both low and high synchrony (**Figure 6C**, low synchrony: control 34 ± 2.3 ms vs. LED stim 38.7 ± 2 ms, n= 7, p= 0.02; high synchrony: control 18.9 ± 2.2 vs. LED stim 24.7 ± 2.6 ms, n= 7, p<0.001). Additionally, the current needed to evoke a spike in MCs, across all sim-EPSPs, was significantly reduced in presence of light stimulation, in agreement with the disinhibitory action of the MCPO GABAergic inputs (**Figure 6D**, low synchrony peak: p<0.001; high synchrony peak: p<0.001). Together, these results indicate that MCPO GABAergic inhibition of GCs results in disinhibition of MCs, which in turn results in a decrease in the firing precision of the output neurons.

**Figure 6.**
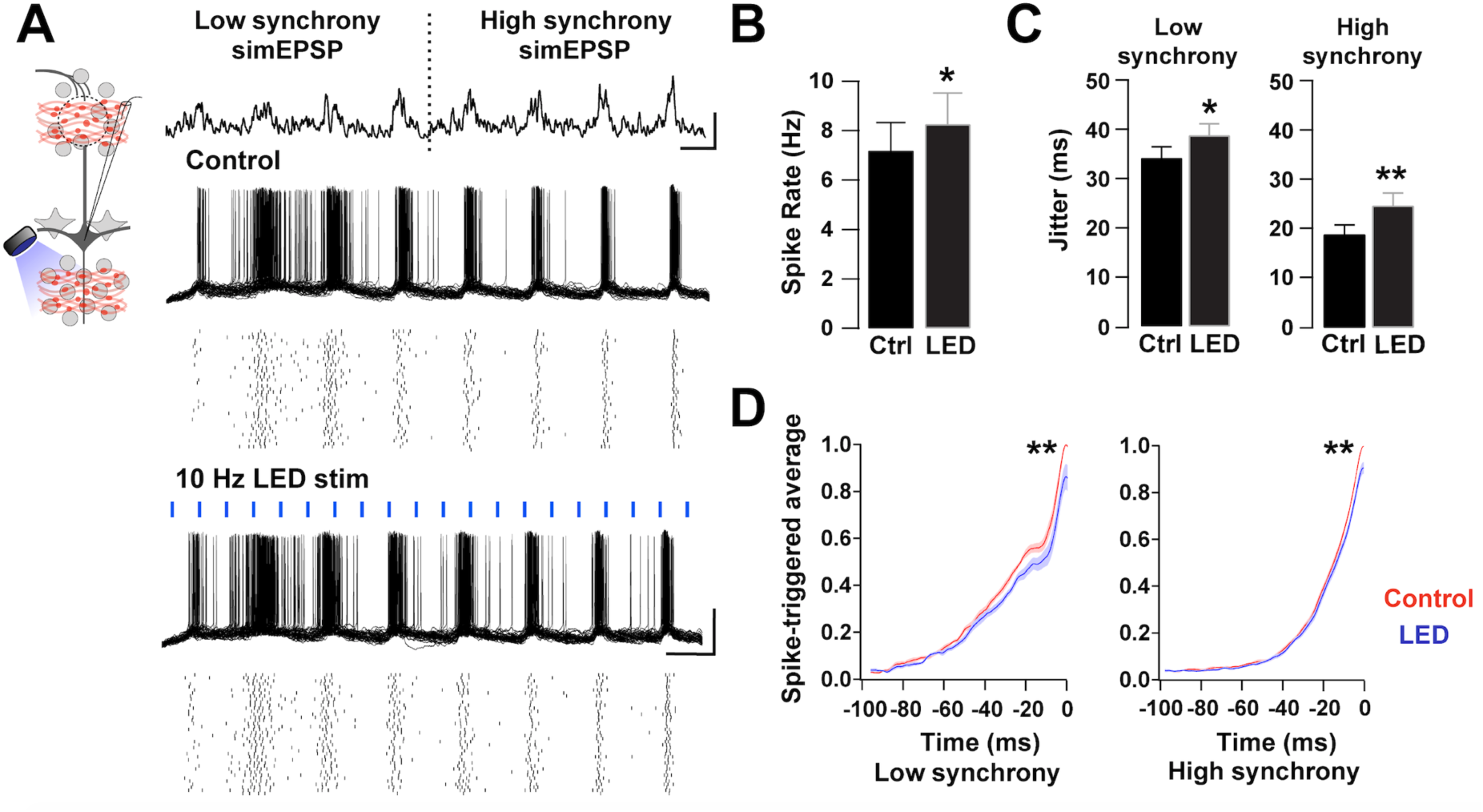
Activation of BF GABAergic inputs desynchronizes MCs. **(A)** Left top, diagram of the experimental configuration; MCs were recorded in current clamp while axons of BF GABAergic neurons expressing ChR2 were locally activated in the GCL by blue light (5 ms, 10 Hz). MCs were stimulated with fluctuating currents that simulate sensory input of increasing synchrony on a 4 Hz sine wave (upper trace). The simulated current injection had low (first 4 current bursts) and high synchrony (last 4 current bursts). Overlaid voltage traces from a MC held at −60 mV in response to the current stimuli, during control (upper traces) and blue light stimulation (LED 5ms, 10 Hz, lower traces). Raster plots underneath show single cell responses in 40 trials. The scale bars are 100 pA and 200 ms (top); 40 mV and 200 ms (bottom). **(B)** The overall firing rate of MCs was significantly increased in the presence of blue light stimulation (n= 7, p= 0.02). **(C)** The spike jitter was significantly increased during low synchrony and high synchrony simulated sensory inputs in the presence of blue light stimulation (low synchrony, p= 0.02; high synchrony, p<0.001). **(D)** Spike-triggered average during low synchrony (left) and high synchrony (right) in the presence (red) or absence (blue) of blue light stimulation. The peak current needed to elicit spikes was smaller in the presence of blue light stimulation (low synchrony peak: p<0.001; high synchrony peak: p<0.001).

## DISCUSSION

The OB receives a rich GABAergic innervation from the MCPO, a region of the BF, and chemogenetic silencing of these neurons impairs olfactory discrimination (Nunez-Parra et al., 2013). Here, we provide new mechanistic insights on how long-range GABAergic inhibition shapes early sensory processing by influencing local inhibition in the OB. BF inhibition directly regulates local inhibitory neurons, including the GCs and PGCs, producing a net disinhibition of the OB output neurons. This disinhibition affected the function of MCs at two levels; in the temporal domain, activation of BF inhibition produced a phasic increase in the firing of MCs and a decrease in their spiking precision. Additionally, activation of LRGNs reduced the extent of dendrodendritic inhibition at GC-MC synapses, suggesting that top-down GABAergic inhibition can also regulate MCs function in the spatial domain. At the circuit level, activation of the GABAergic feedback produced a specific modulation of inhibition across the glomerular and GC layers. Phasic activation of BF-LRGNs resulted in a differential modulation of the intensity of *θ* and *γ* band oscillations across these two layers. Thus, phasic activation of BF long-range GABAergic inhibition is poised to influence both the spatial and temporal aspects of early olfactory processing.

The BF contains a heterogeneous population of projection neurons that innervate cortical and subcortical areas, where they have an important role in the state-dependent regulation of sensory circuits (Jourdain et al., 1989; Zaborszky et al., 2012; Yang et al., 2014; Hangya et al., 2015; Zant et al., 2016). The MCPO is the most important source of GABAergic projections to the OB, however the function of these inhibitory neurons has been difficult to assess due to the presence of other cell types in the BF, including cholinergic and glutamatergic neurons (Zaborszky et al., 2012; Yang et al., 2017). Our functional and neuroanatomical studies provide direct evidence that these GABAergic projections use GABA as a main transmitter. The cholinergic marker ChAT was not present in MCPO GABAergic neurons and the fast-inhibitory currents elicited by their activation were insensitive to cholinergic antagonists. Thus, although we cannot rule out the possibility that ChAT is expressed at low levels in OB projecting MCPO neurons, undetected by our immunoassay, or that MCPO Gad2+ neurons can release other neurotransmitters (Trudeau and El Mestikawy, 2018), our evidence supports a main GABAergic phenotype for these neurons. Phasic activation of MCPO GABAergic neurons produce a fast inhibition in local OB inhibitory neurons, which distinguish them from a different subtype of BF projection neurons previously described (Saunders et al., 2015; Case et al., 2017).

Functionally, fast MCPO GABAergic inhibition shapes the OB output by regulating local inhibitory circuits, instead of directly acting on the output neurons. LRGNs preferentially elicit GABAergic currents on inhibitory neurons in both MOB and AOB. Although, it is possible that small responses could exist in the output neurons, the absence of responses in these neurons is also consistent with neuroanatomical studies indicating that BF GABAergic afferents only target inhibitory neurons in the OB (Gracia-Llanes et al., 2010). This bias towards GABAergic targets has been reported for other long-range inhibitory projections in the brain (Freund and Antal, 1988; Gulyás et al., 1990, 1991; Freund and Gulyás, 1991; Martínez-Guijarro and Freund, 1992; Melzer et al., 2012; Caputi et al., 2013; Gonzalez-Sulser et al., 2014). The function of this biased pattern is unknown, however, given the essential participation of inhibitory circuits in network synchronization (Buzsáki and Chrobak, 1995), it has been proposed that long-range GABAergic inhibition modulates temporal dynamics in target circuits (Hangya et al., 2009; Kim et al., 2015; Viney et al., 2018). At the network level, fast feedforward inhibition of local GABAergic neurons by BF-LRGNs decreases the intensity of evoked *θ* and *γ* band oscillations in the OB, through direct activation of GABA_A_R in a circuit specific manner. Oscillations are inherent to olfaction (Kay et al., 2009), and underlie fine odor discrimination and high cognitive tasks (Stopfer et al., 1997; Beshel et al., 2007; Nusser et al., 2013). Interestingly, disruption of GABA_A_R in GCs increases *γ* oscillations (Nusser et al., 2013), further supporting the possibility that inhibition of GCs influences synchronized activity in the OB. Thus, we propose that BF-LRGNs function as a phasic feedback mechanism that modulates synchronous activity in the OB, thus avoiding hypersynchrony (Khazipov, 2016). A similar mechanism has been proposed in the thalamus, where reduction in the GABA_A_R mediated inhibition intensifies thalamocortical oscillatory activity (Huntsman et al., 1999). Interestingly, studies *in vivo* have shown that cortically projecting BF GABAergic neurons increase *γ* band oscillations by modulating local fast spiking (FS) inhibitory neurons (Kim et al., 2015). Thus, the EPL-I neurons in the MOB could have a similar function as they also exhibited fast inhibition upon MCPO GABAergic activation. Future experiments should evaluate the contribution of FS neurons to *γ* oscillations in the MOB and their regulation by BF inhibition. Nevertheless, these changes in synchrony at the network level can also be explained by decorrelation of activity in the output neurons, as BF-LRGN activation reduced spike precision on MCs in response to a simulated sensory input. It is noteworthy that GCs have been proposed to participate in the generation of highly precise firing in MCs (Schoppa, 2006), which is thought to underlie temporal encoding in the OB (Kepecs et al., 2006; Shusterman et al., 2011).

Since different odor molecules activate overlapping subpopulations of mitral/tufted cells (M/TCs), pattern separation is an essential aspect of olfactory processing. Lateral inhibition via dendrodendritic inhibition is thought to provide a mechanism for this tuning specificity across M/TC subpopulations (Yokoi et al., 1995; Isaacson and Strowbridge, 1998; Mori et al., 1999; Christie et al., 2001; Shepherd, 2004). Thus, top-down GABAergic inhibition of local circuits can influence the M/TCs interactions in the spatial domain and modulate tuning specificity of M/TCs. BF GABAergic inhibition directed either to the glomerular or the GCL circuits greatly reduced the extent of inhibition in MCs. Interestingly, the density of innervation by MCPO GABAergic axons is highest in the GCL supporting a strong influence on GCs. This is in agreement with studies that suggest that odor pattern separation partially depends on GABAergic inhibition of GCs (Abraham et al., 2010; Friedrich and Wiechert, 2014; Gschwend et al., 2015), and that chemogenetic silencing of MCPO LRGNs impairs the discrimination of similar odors (Nunez-Parra et al., 2013). On the other hand, in the glomerular domain, the decay of the IPSC elicited by MCPO axons activation in PGCs is significantly faster than for GCs. PGCs are reciprocally connected with M/TCs, from which they receive a strong excitation (Murphy et al., 2005). Consistent with our findings, a recent report described robust IPSCs in a subpopulation of PGCs elicited by activation of BF GABAergic neurons (Sanz Diez et al., 2019). This dendrodendritic interaction is thought to gate the glomerular output by regulating the activity of M/TCs (Wachowiak and Shipley, 2006; Gire and Schoppa, 2009; Shao et al., 2012), suggesting that phasic activation of BF-LRGNs can rapidly modulate the glomerular circuits, strongly impacting the strength of the incoming sensory input. Furthermore, MCs receive synchronous GABAergic inputs from GCs in an activity-dependent manner, which facilitates the integration of odor information via lateral interactions among MCs (Arnson and Strowbridge, 2017). Similarly, fast and synchronous inhibition by BF-LRGNs could control a large spatial domain, through a temporally precise GABA release onto the local OB inhibitory network. Together these results suggest that the BF GABAergic input to the OB is well suited to rapidly modulate the extent of local inhibition in the glomerulus and the lateral dendrites of MCs, and thus modulate odor processing.

Our results underscore the view that GCs integrate inhibition from two sources: top-down inhibition from MCPO afferents and inhibition from local interneurons (Pressler and Strowbridge, 2006; Eyre et al., 2008; Burton and Urban, 2015), however these sources of inhibition may play different functions on GCs. The timing of GABAergic inhibition is shaped by the kinetics of the IPSC, which strongly depends on the type of GABA_A_R expressed postsynaptically and the mechanisms of GABA release (Okada et al., 2000; Ortinski et al., 2004; Succol et al., 2012). Phasic activation of MCPO GABAergic inputs elicited a fast synchronized release of GABA, suggesting a tight coupling between presynaptic action potentials and the release events (Kaeser and Regehr, 2014). However, the IPSC elicited by MCPO inputs in GCs had a slower time course (∼40 ms) compared to the decay of the IPSC elicited by local inputs (∼6 ms) (Eyre et al., 2012), suggesting that these sources of inhibition may have a different function in the temporal domain. In the hippocampus IPSCs with fast decay kinetics are thought to facilitate *γ* oscillations, while IPSCs with slow decays likely control postsynaptic excitability (Bartos et al., 2002). Thus, it is possible that the slower decay of top-down inhibition has a stronger influence in the excitability of GCs, and therefore is more suited to modulate OB spatiotemporal dynamics by influencing MCs firing. Furthermore, the fast rise time of the min-IPSC suggested a predominant perisomatic targeting of the MCPO GABAergic input to GCs, perhaps reflecting a convergent regulation by top-down inputs. Thus, BF inhibition could have a strong influence on the excitatory inputs that also target the proximal region of the GCs (Balu et al., 2007). A fast disinhibitory function of BF-LRGNs is especially relevant, since all the afference that the OB receives from the olfactory cortices is excitatory on the local interneurons, and produces an overall inhibition of the OB output (Boyd et al., 2012, 2015; Markopoulos et al., 2012; Rothermel and Wachowiak, 2014; Otazu et al., 2015).Thus, somatic inhibition of GCs could reduce the likelihood of the excitatory feedback in generating somatic action potentials (Freund and Katona, 2007), limiting the effect of the cortical feedback on modulating M/TCs, further experiments are needed to determine the temporal widow in which this regulation can occur.

Interestingly, in *in vivo* recordings from OB projecting MCPO neurons, these cells can be directly excited by activation of the piriform and entorhinal cortices (Paolini and McKenzie, 1997), suggesting modulation of MCPO LRGN activity by incoming odor-elicited activity from olfactory areas. Since GCs are targeted by the MCPO GABAergic axons, these projections could participate in an OB-BF feedback loop that can rapidly modulate the temporal code in the OB. The highly branched BF-LRGN innervation across the OB cellular layers in addition to a relatively small number of OB projecting neurons in the MCPO (∼680 GABAergic neurons) (Gracia-Llanes et al., 2010), suggests a broad postsynaptic control by the GABAergic axons. Single axons could influence a large number of interneurons in the OB, thus contributing to spatial code modulation. Thus, we hypothesize that top-down inhibition provides a rapid disinhibitory feedback to ongoing odor-induced activity in the OB, influencing the temporal and spatial dynamics of odor coding in the OB, including gain control and tuning specificity of the output neurons.

## METHODS

### Animals

All experiments were conducted following the guidelines of the Institutional Animal Care and Use Committee of the University of Maryland, College Park. For our experiments we used wild type C57BL/6 (JAX, stock #664) and Gad2-IRES-Cre mice (JAX, stock #010802) of both sexes, ranging in age from one to four months, from breeding pairs housed in our animal facility.

### Stereotaxic injections

Deep anesthesia of Gad2-Cre mice was induced with 2% isoflurane at a rate of 1 L/min and adjusted (1-1.5%) over the course of the surgery. Body temperature was maintained using a heating pad. An intraperitoneal injection of carprofen (5 mg/Kg) was used as analgesic and a solution of povidone-iodine (Betadine) as antiseptic. During the surgery, eyes were lubricated using a petrolatum ophthalmic ointment (Paralube). GABAergic projection neurons in the basal forebrain were retrogradely labeled using a unilateral injection of AAVrg-hSyn-DIO-eGFP in the OB (50 nL, Catalog #50457-AAVrg, Addgene), guided with a stereotaxic apparatus (Kopf, Catalog #940), and using the following stereotaxic coordinates (mm): −D/V 0.4, ±M/L 0.8, A/P +6. This retrograde injection in the OB sparsely labeled neurons in the anterior olfactory nucleus, as recently shown (Hanson et al., 2020). To express channelrhodopsin-2 (ChR2) in LRGNs, Gad2-Cre mice were bilaterally injected with AAV5-CAG-Flex-ChR2-tdTomato (200 nL, Catalog #18917, Addgene) in the MCPO region of the BF using the following stereotaxic coordinates (mm): D/V −5.4, M/L ±1.63, A/P +0.14. For histology experiments, the control virus AAV5-CAG-Flex-tdTomato (200 nL, Addgene) was used to anterogradely label MCPO GABAergic axons. For both retrograde and anterograde labelling of LRGNs, electrophysiological or histological experiments were conducted 3 weeks, or later, post-surgery.

### Confocal imaging and immunofluorescence

To directly visualize the expression of the reporter gene (tdTomato or eGFP), mice were transcardially perfused with cold 4% PFA diluted in 0.1 M PBS, pH 7.4. Brains were then harvested and post fixed overnight at 4°C in the same fixative. Brain tissue was sliced in sections of 50 μm on a vibratome, the nuclei stained with DAPI (1:1500, Catalog #D1306, Invitrogen) and mounted in a solution of Mowiol-DABCO. Mowiol mounting media was made in batches of 25 mL containing 9.6% w/v mowiol (Catalog #475904, Millipore), glycerol 24% w/v, 0.2 M Tris (pH 6.8), 2.5% w/v DABCO (antifade reagent, Catalog #D2522, Sigma) and Milli-Q water. For immunofluorescence experiments, free floating brain sections (50 μm) were first blocked with donkey serum (10%, Catalog #S30-M, Millipore) in PBS supplemented with Triton X-100 (0.1% v/v, Catalog #T8787, Millipore, PBS-T) for 1 h at room temperature to block unspecific biding sites. Samples were then incubated overnight at 4°C with a goat primary antibody anti-ChAT (1:500, Catalog #AB144, Millipore) and 2.5% donkey serum in PBS-T with gentle rocking. The primary antibody was then washed with PBS-T for at least 30 min before incubation with a donkey anti-goat antibody coupled to Alexa-647 (1:750, Catalog #A-21447, Invitrogen). Finally, slices were stained with DAPI, dried and mounted using Mowiol-DABCO. Control sections not exposed to the primary antibody were devoid of immunostaining and were used to set background values on the microscope. Images were acquired using a Leica SP5X confocal microscope, with appropriate brightness and contrast adjustments, and immunostained cells counted blindly using ImageJ (NIH).

### Whole cell recordings

Patch clamp recordings in brain slices were conducted as previously described (Nunez-Parra et al., 2013) using a dual EPC10 amplifier (HEKA). Briefly, we used a vibratome (VT1000S, Leica) to obtain horizontal 250 μm slices. Sectioning was done using a cold low Ca^2+^ (0.5 mM) and high Mg^2+^ (3 mM) artificial cerebrospinal fluid solution (ACSF). Slices were then placed in normal Ca^2+^ and Mg^2+^ ACSF and left to recover for 30-45 min at 37°C. The normal ACSF had the following composition (in mM); 125 NaCl, 25 NaHCO_3_, 1.25 NaH_2_PO_4_, 3 KCl, 2 CaCl_2_, 1 MgCl_2_, 3 myo-inositol, 0.3 ascorbic acid, 2 Na-pyruvate and 15 glucose, and it was continuously oxygenated with 95% O_2_ and 5% CO_2_. After recovery, slices were transferred to a recording chamber on an Olympus BX51W1 DIC microscope. Neurons were visualized using 4x and 40x objectives (LUMPlanFI/IR, Olympus). The evoked inhibitory postsynaptic currents (eIPSCs) were recorded at a holding potential of 0 mV using an internal solution with the following composition (in mM): 125 Cs-gluconate, 4 NaCl, 2 MgCl_2_, 2 CaCl_2_, 10 EGTA, 10 HEPES, 2 Na-ATP, 4 Mg-ATP and 0.3 GTP. Alternatively, the eIPSCs were recorded at −70 mV using an internal solution of the following composition (in mM): 150 CsCl, 4.6 MgCl_2_, 0.1 CaCl_2_, 10 HEPES, 0.2 EGTA, 4 Na-ATP and 0.4 Na-GTP. The pH of internal solutions was adjusted to pH 7.3 with CsOH. In some experiments, CaCl_2_ was replaced by equimolar amounts of SrCl_2_ in the ACSF. No Ca^2+^ chelators were added to this solution. To confirm the identity of the recorded neurons, and morphological reconstruction, the fluorophore Alexa-594 (20 µM, Invitrogen) was included in the internal solution in a subset of experiments. Post-recording filled neurons were fixed overnight at 4°C in PFA 4% and mounted with Mowiol-DABCO. Neurons were imaged under a confocal microscope and reconstructed using Neurolucida (MBF Bioscience) or neuTube (Feng et al., 2015). Recording were performed at room temperature (21°C). Patch pipettes were pulled using a horizontal puller (P-97, Sutter Instrument) from thick wall borosilicate glass capillaries (Sutter Instrument), having a resistance of ∼3-6 MOhm. All chemicals were obtained from Sigma Aldrich. Drugs were prepared from stocks stored at −20°C and diluted into ACSF; gabazine (Catalog #1262, Tocris), mecamylamine hydrochloride (Catalog #2843/10, Tocris), atropine (Catalog #A0132, Millipore-Sigma), kynurenic acid sodium salt (Catalog #3694, Tocris).

### LFP recordings and optogenetic stimulation

Local field potentials in the OB were recorded in 250 μm brain slices using 200-300 KOhm glass electrodes filled with ACSF. To induce oscillations in the OB, a brief stimulation (100 µA, 100 Hz during 50 ms) was delivered to the ON using a stimulus isolation unit (ISO-Flex, A.M.P.I) controlled by the amplifier. OSN axon bundles were readily seen under DIC optic. For optogenetic stimulation of the GL or GCL a collimated LED (473 nm, Thor Labs) was used to deliver brief light pulses through a 40x objective focused on either layer (which were at least ∼400 µm apart), controlled by a TTL pulse triggered by amplifier. The intensity of the light beam was adjusted depending of the level of ChR2 expression from 1-3 mW mm^−2^, measured after the objective. Trials were alternated between control and optogenetic stimulation conditions.

### Data analysis

Electrophysiology data was analyzed using the IgorPro (WaveMetrics) and MATLAB (MathWorks) software. Only events with fast current kinetics were included in the analyses (rise time <4 ms and decay time <100 ms). For the quantification of the currents elicited by light stimulation, we calculated the transferred charge by integrating the current within a 300 ms window following the end of the light pulse and subtracting the baseline charge before the light pulse. The decay time was measured by fitting a double exponential decay function to the current relaxation and computing the weighted time constant (τ_w_) as τ_w_ = (a_1_τ_1_ + a_2_τ_2_)/ (a_1_ + a_2_), where a and τ are the amplitude and time constant of the first (1) and second (2) exponentials, respectively. The rise time was estimated by measuring the time elapsed from 10 to 90% of the current peak amplitude. The onset time was measured by fitting a sigmoid function from stimulus onset to the response peak and then computing the maximum curvature point by solving the 4^th^ derivative of the fitted curve set equal to zero (Fedchyshyn and Wang, 2007). Spike jitter was measured as total standard deviation of the timing of the action potential peaks (Mainen and Sejnowski, 1995; Gutkin et al., 2003). Spike-triggered averages were calculated using an average of the current stimuli corresponding to the 100 ms prior to each action potential. For LFP analysis, stimulus artifacts were digitally removed, and traces were filtered with a 2^nd^ order 300 Hz Butterworth low-pass filter. Spectral analysis was then conducted using the Chronux toolbox (http://www.chronux.org) using a multitaper spectral estimation (Bokil et al., 2010). For the histograms of axonal density, the cellular layers of different field of views were aligned horizontally using the nuclear staining as a guide. Mean pixel intensity values were computed across the horizontal axis and normalized to the overall maximal intensity value. Data is shown as the mean ± S.E.M, unless otherwise specified. Statistical analysis was done using a two-tailed t-test and significance was set at p<0.05 (*= p<0.05, **= p<0.01). Statistical power was evaluated using G*Power (Faul et al., 2009).

## AUTHOR CONTRIBUTIONS

P.S.V. and R.C.A. designed research; P.S.V. and R.H. performed research; P.S.V. and R.H. analyzed data; PV and R.C.A. wrote the paper.

## EXTENDED FIGURES

**Figure S1.**
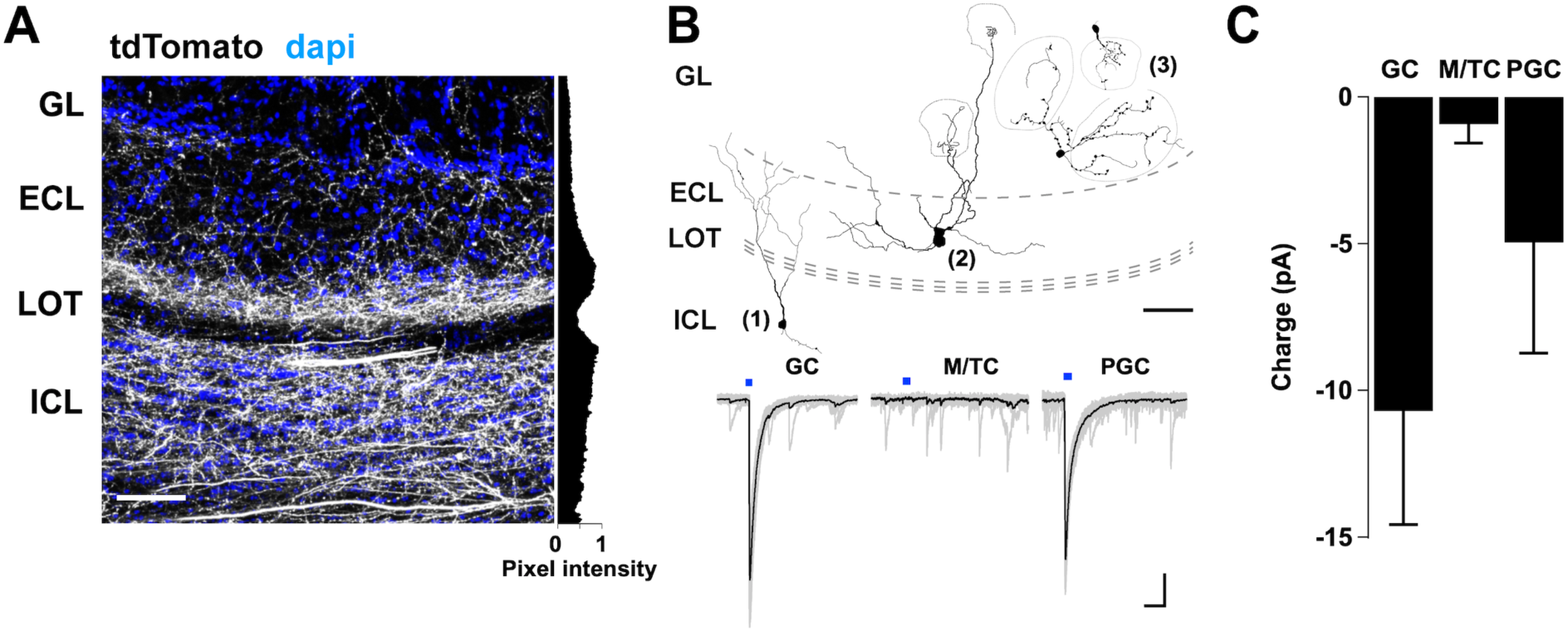
Inhibitory neurons are postsynaptic partners of MCPO long-range GABAergic neurons in the AOB. **(A)** Confocal image of a sagittal section of the accessory OB (AOB) expressing tdTomato in the MCPO GABAergic axons (shown in white). Nuclei stained with dapi are shown in blue and delineate the different cellular layers of the AOB. The histogram on the right shows the average normalized pixel intensity across the cellular layers; GL, 0.29 ± 0.05; ECL, 0.62 ± 0.11 and GCL, 0.92 ± 0.06 (n= 4). The scale bar is 100 μm. **(B)** Upper panel, example of reconstructed neurons, post-recording; 1, GC; 2, mitral/tufted cell (M/TC); 3, PGC. The morphology of the neurons was reconstructed from confocal images of fixed cells that were filled with Alexa Fluor-594 during the recordings. The scale bar is 100 μm. Bottom, sample eIPSCs elicited by LED stimulation (5 ms) of GABAergic axons expressing ChR2. GABAergic currents were observed in GCs and PGCs, but not in the output neurons, the M/TCs, in recordings at −70 mV. GL, glomerular layer; ECL, external cellular layer; LOT, lateral olfactory tract; ICL, internal cellular layer. The scale bar is 200 ms, and 50 pA **(C)** Bar graph showing the total charge transferred during the GABAergic eIPSCs in distinct cell types in the AOB; responses are observed in the main inhibitory types, but not in the output neurons (GC, n= 12; M/TC, n= 6, PGC, n= 5).

